# miRVim: Three-dimensional miRNA Structure Data Server

**DOI:** 10.1101/2023.11.29.569154

**Authors:** Vishal Kumar Sahu, Ankita Subhadarsani Parida, Amit Ranjan, Soumya Basu

## Abstract

MicroRNAs (miRNAs), a distinct category of non-coding RNAs, exert multifaceted regulatory functions in a variety of organisms, including humans, animals, and plants. The inventory of identified miRNAs stands at approximately 60,000 among all species and 1,926 in *Homo sapiens* manifests miRNA expression. Their theranostic role has been explored by researchers over the last few decades, positioning them as prominent therapeutic targets as our understanding of RNA targeting advances. However, the limited availability of experimentally determined miRNA structures has constrained drug discovery efforts relying on virtual screening or computational methods, including machine learning. To address this limitation, miRVim has been developed, providing a repository of human miRNA structures derived from both two-dimensional (MXFold2, CentroidFold, and RNAFold) and three-dimensional (RNAComposer and 3dRNA) structure prediction algorithms, in addition to experimentally available structures from the RCSB PDB repository. This data server aims to facilitate computational data analysis for drug discovery, opening new avenues for advancing technologies such as machine learning-based predictions in the field. The publicly accessible structures provided by miRVim, available at https://mirna.in/miRVim, offer a valuable resource for the research community, advancing the field of miRNA-related computational analysis and drug discovery.

## Introduction

A subclass of non-coding RNAs (ncRNA), microRNAs (miRNAs) are small molecules that have been recognised as an important regulator of gene expression in various biological processes. Two independent researchers in 1993 identified that the *Caenorhabditis elegans* heterochronic gene lin-4, a small noncoding RNA, plays an important role in post transcriptional regulation in temporal pattern formation (Wightman et al., 1993). After a seven-year hiatus, it was empirically demonstrated that let-7, heterochronic gene in *C. elegans*, in conjunction with lin-4 RNA possesses the capacity to initiate the temporal cascade of regulatory heterochronic genes (Reinhart et al., 2000). Subsequently, this discovery catalysed extensive research endeavours substantiating the presence of a substantial cohort of small ncRNAs endowed with regulatory functions, which were later termed as miRNAs (Lagos-Quintana et al., 2001).

The miRNAs are involved in the regulation of gene expression by regulating gene transcription, affecting mRNA splicing, directing DNA methylation, or by preventing RNA export (MacFarlane & Murphy, 2010). Due to their ability to fine-tune gene expression, miRNAs have garnered significant attention as potential therapeutic targets and biomarkers for various diseases. As an example miR-17/92 cluster has been proven to be important in cell cycle, proliferation, apoptosis and other pivotal processes. The cluster has been associated with normal development and it has been the first group of miRNAs to be implicated in Feingold syndrome (Mogilyansky & Rigoutsos, 2013). The miRNA-92a and -21 have been reported to be circulating and minimally invasive biomarkers in primary breast cancer (Si et al., 2013). In myasthenia gravis, miR155 has been demonstrated to be required for antibody production after vaccination with attenuated Salmonella and posting miR-155 inhibitor as a promising approach for the clinical therapy (Y.-Z. Wang et al., 2014).

Comprehending the structural attributes of miRNAs assumes paramount significance in elucidating their functional mechanisms and probing their ramifications across diverse biological *milieus*. Current estimates predicate that miRNAs wield regulatory control over approximately 60% of human genes (Bartel, 2009). Their involvement extends to a panoply of biological processes, encompassing cell cycle modulation (Carleton et al., 2007), apoptosis (Jovanovic & Hengartner, 2006), metabolic regulation (Boehm & Slack, 2006), and the orchestration of developmental and differentiation (Harfe, 2005). Furthermore, miRNAs have been unequivocally implicated in an array of maladies, including neurodegenerative and metabolic abnormalities (Poy et al., 2004; Rottiers & Näär, 2012), various forms of cancer (Dalmay & Edwards, 2006), as well as clinically important diseases encompassing autoimmune diseases to myocardial infarction (Soifer et al., 2007).

The miRNAs are essential for controlling gene expression in almost all stages of cancer. Dysregulated miRNA profiles are associated with tumour formation, angiogenesis, progression, metastasis, and therapy resistance (Acunzo et al., 2015; Carleton et al., 2007; Feng et al., 2011; Li et al., 2018; Q. Wang et al., 2023; Yekta et al., 2004). The ability of miRNA signatures to distinguish between healthy and cancerous tissue as well as many subtypes of a given tumour has been demonstrated. Additionally, it has been demonstrated that they serve as biomarkers for early diagnosis and fundamentally contribute to therapy resistance (Calin & Croce, 2006; Dalmay & Edwards, 2006; Iorio & Croce, 2012; Wilk & Braun, 2018). MiRNA studies have explored deeply into their role in drug resistance in recent years (He et al., 2020).

The stability and binding selectivity of miRNAs are determined by the spatial occupancy of atoms participating in the structure formation showing their structural significance equivalent to the protein molecules (Reddy, 2015). The structural information of these molecules is exploited for drug discovery, pattern recognition, and multiple other purposes like stability of complexes. The molecular docking (Rath et al., 2016) and molecular dynamics simulation studies (Y. Wang et al., 2010) require clear understanding of three-dimensional structural complexity for the precise estimation of intermolecular interactions. There are about 1 million experimentally determined unique structures of proteins since 1971. The problem of protein structure determination with atomic accuracy has recently been solved through AlphaFold2 (Jumper et al., 2021) supporting and accelerating the drug discovery by availing more than 200 million protein structures through AlphaFold Protein Structure Database (Varadi et al., 2022). The AlphaFold database has contributed significantly as the same would take enormous time and effort to determine remaining structures.

Understanding the molecular processes underlying miRNA function is costly using *in-vitro* methods as well as limitations using *in-silico* methods due to the scarcity of experimentally identified miRNA structures. Multiple databases housing experimentally proven interactions or binding with highly sophisticated methods or predictive methods are available like miRTarBase (Huang et al., 2021), CLIPdb (Yang et al., 2015), or miRDB (Chen & Wang, 2020), however atomistic levels of interaction may not be available. There are multiple proteins like AGO2, TNRC6B, CNBP, or TNRC6A, have been already established to be interacting with miRNA supported by miRTarbase and CLIPdb published data, however targeting or deciphering exact interaction might require further in vitro studies which may be expensive and take additional time to validate the interactions delaying drug discovery.

The advancement of computational techniques for predicting miRNA-target interactions and designing effective miRNA-based therapies is impeded by the absence of a comprehensive miRNA structural database. The enhanced understanding of miRNA biology and the discovery of novel therapeutic approaches depend on efforts to address structural characterization of miRNAs and development of high-throughput experimental methods for their structure determination (Riolo et al., 2021).

In the view of scarcely available structures of microRNAs, the miRVim was developed as a data server that contains predicted structure of human miRNAs curated using open-source tools and established 2D and 3D RNA structure prediction techniques. This was done to overcome the limitation of miRNA structures leading to drug screening targeting miRNAs and machine learning models utilising the 3D structures of miRNAs.

## Results

### Statistics of available structures

The miRVim contains miRNA sequences from miRBase and their corresponding 2D and 3D structures. The length of 1,729 precursor miRNAs available in miRBase ranged between 41-180 nt while for 2,924 mature miRNAs it was 16-28 nt in the species *Homo sapiens*. The miRVim database includes a total of 4,653 miRNA sequences, including precursor- and mature-miRNAs. The database includes 13,971 2D structures of miRNAs, determined using RNAFold, MXFold2 and CentroidFold algorithms using their respective methods. Furthermore, the database contains 17,045 3D structures of miRNAs, predicted using 3dRNA and RNAComposer server (Table 1). However, only 56 experimentally determined structures are available.

**Table 1:**
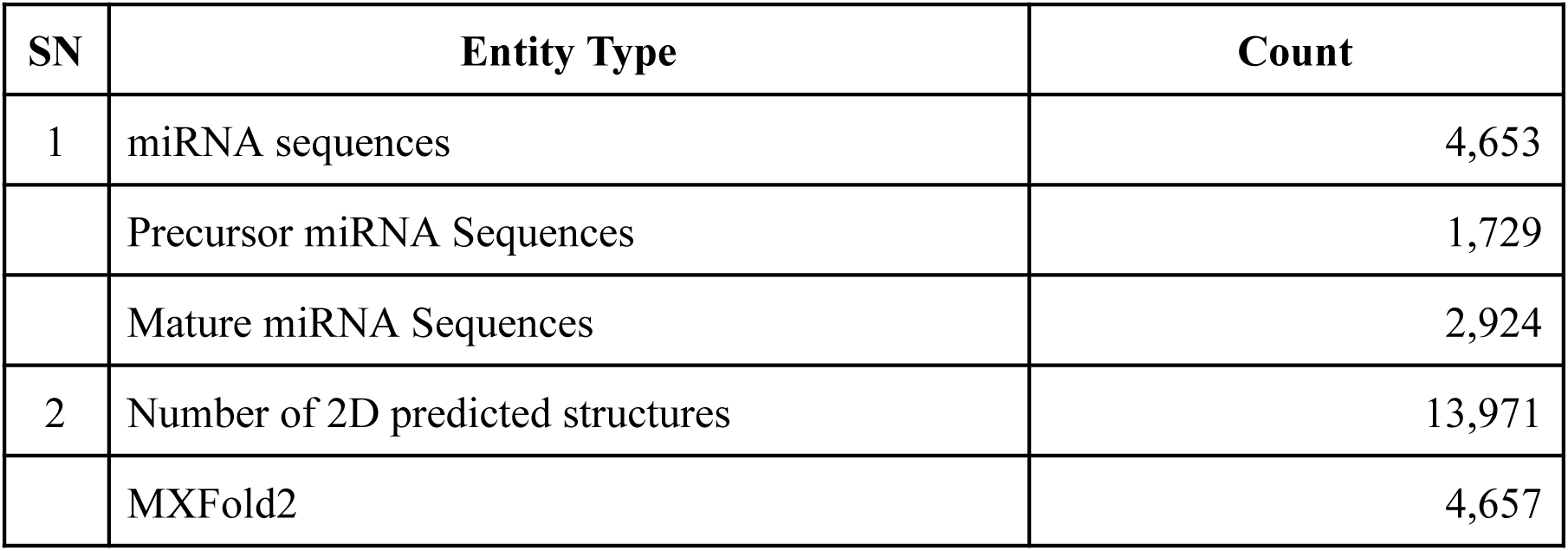
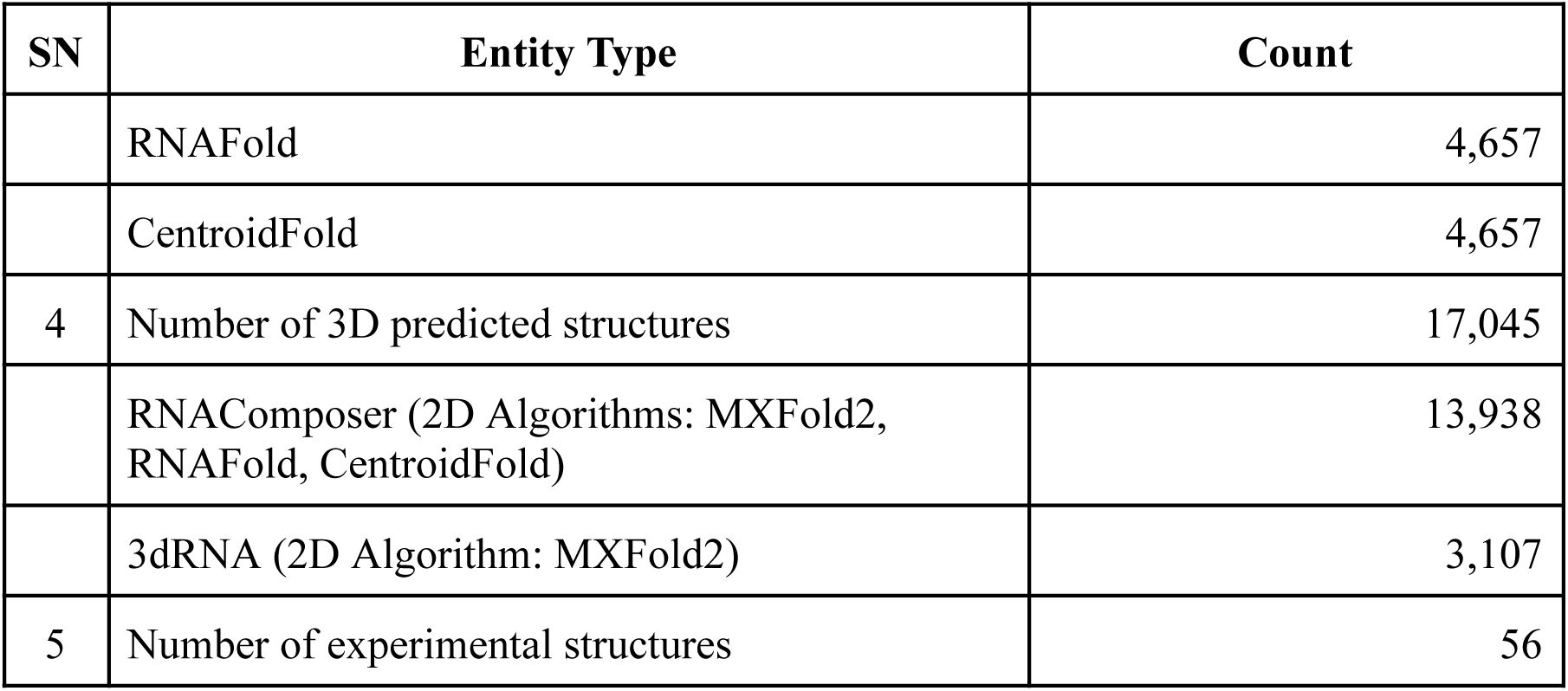
Statistics of available structures in the miRVim server.

| SN | Entity Type | Count |
| --- | --- | --- |
| 1 | miRNA sequences | 4,653 |
|  | Precursor miRNA Sequences | 1,729 |
|  | Mature miRNA Sequences | 2,924 |
| 2 | Number of 2D predicted structures | 13,971 |
|  | MXFold2 | 4,657 |
|  | RNAFold | 4,657 |
|  | CentroidFold | 4,657 |
| 4 | Number of 3D predicted structures | 17,045 |
|  | RNAComposer (2D Algorithms: MXFold2, RNAFold, CentroidFold) | 13,938 |
|  | 3dRNA (2D Algorithm: MXFold2) | 3,107 |
| 5 | Number of experimental structures | 56 |

### Web interface and data availability

The web interface of miRVim accepts partial or complete input query for the name of miRNA, identifier, gene name, or short sequence (Figure 1a) through the search field. Results matching the query are returned (Figure 1b) requiring further action. Individual results containing detailed results about the miRNA entity can be expanded to explore, visualise, and download related structures and details (Figure 1c and Figure 1d).

**Figure 1:**
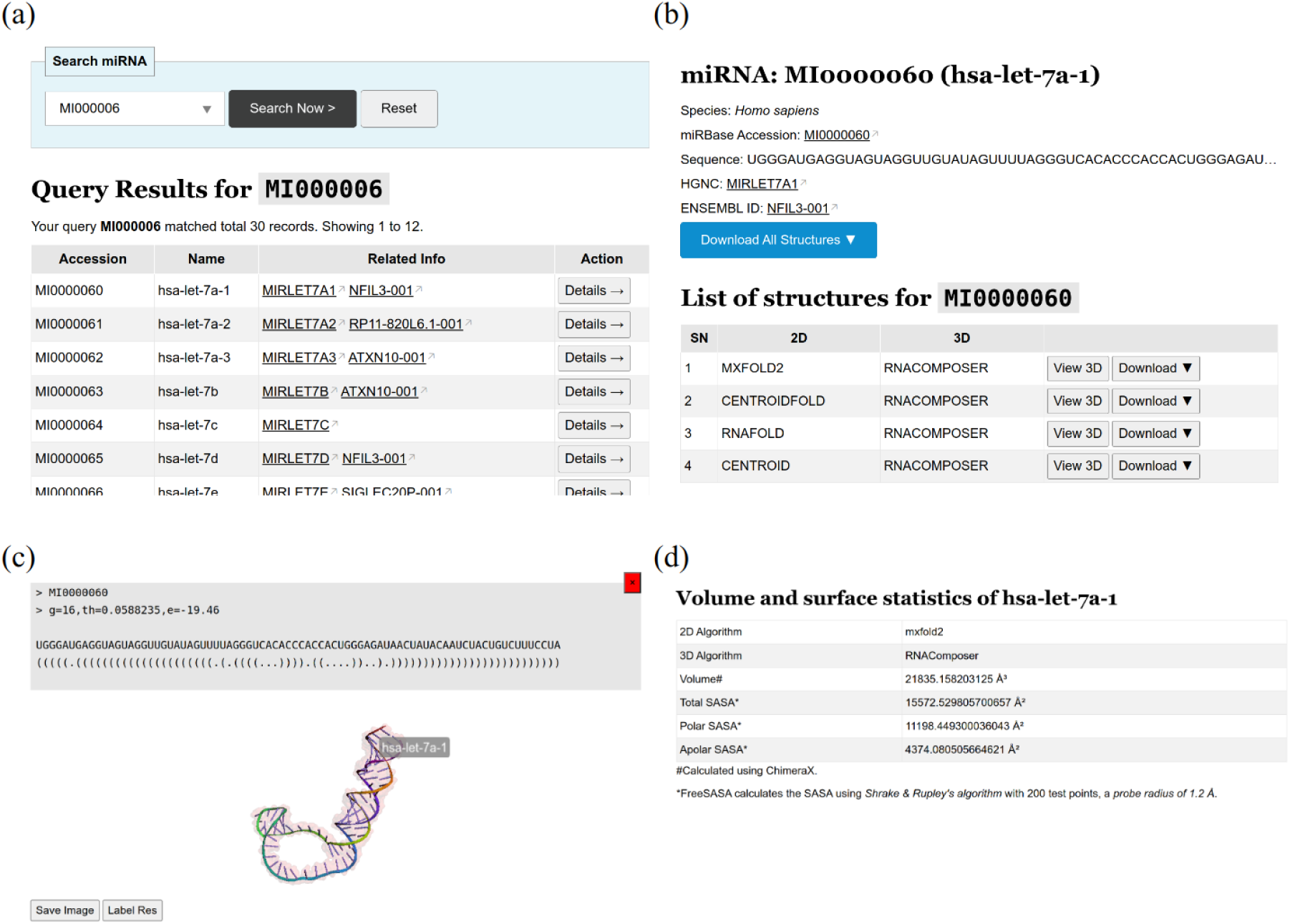
Web UI search box to query miRNA. It finds results by partial or complete match of accession number, sequence, gene name, or transcript name of miRNAs. (a) Multiple records matching the query term, (b) details of selected record from previous action including visualisation and download option, (c) visualisation of three-dimensional structure of the selected miRNA, and (d) volume and solvent accessible surface area statistics along with secondary structure information are available through miRVim data server.

Overall, this database represents a valuable resource for researchers interested in miRNA biology and their interactions with other molecules. The availability of both 2D and 3D structures for a large number of miRNAs allows for a more detailed understanding of miRNA function and regulation using computational methods through structure analysis. The approach may lead to the development of new therapeutic strategies for various diseases from the field of RNA biology.

## Discussion and conclusion

Drug discovery methods involving *in silico* or *in vitro* approach has been primarily focused on the protein molecule and now the trend is shifting towards nucleic acids (Bernetti et al., 2022; Conti & Oppikofer, 2022). The *in silico* study depends on the various attributes of the molecules like sequence, chemical composition, or 3D structure. For proteins, the AlphaFold Database has filled the gap by accelerating the determination of more than 200 million structures. Certainly it has provided enormous support to drug discovery and therapeutics involving protein structures. However, miRNA structure determination and unavailability of a database to provide miRNA structure is also required to support the drug discovery related to miRNAs. The study related to modifications in RNA structure or functional groups, like pseudouridine (Ψ) (Schaefer et al., 2017) or glycosylation (Tyagi et al., 2023), would certainly lead to more complex interactions requiring different approaches for the deduction of three dimensional structures.

Experimental methods verify how miRNA interacts with its targets; however determination of key interactions require exhaustive iterations due to complexity of interacting species using various laboratory techniques involving miRNA-miRNA, miRNA-DNA, or miRNA-protein interactions (Gan & Gunsalus, 2013; Krol et al., 2010; Negoita et al., 2017; Plotnikova et al., 2019). Molecular docking or machine learning approach may precisely deliver the key interactions among the interacting biomolecules paving path to drug discovery targeting those interactions. The three-dimensional structure information is at par with the requirement of discovering key interactions among miRNAs or intermolecular interactions. The services, information, and data provided by the miRVim data server as a unified repository of predicted and experimentally derived structures availed from RCSB PDB. The database serves as a valuable resource addressing the existing gap in openly accessible miRNA structure databases. Its repository of readily accessible miRNA structures offers an expedited avenue for advancing conventional drug discovery methodologies that leverage miRNA structures, notably through the integration of bioinformatics and machine learning techniques.

This comprehensive miRNA structure information provided by initial version of miRVim holds significant utility in various computational endeavours, including molecular docking and dynamics simulations, machine learning-based tools, and *in-silico* pharmacology investigations aimed at elucidating miRNA interactions with other molecular entities. Such insights have the potential to facilitate the identification of promising drug targets, fine tune gene regulation controlling disease progression, and the innovative development of pharmacological agents designed to target miRNAs.

## Methods

### Primary sequence from miRBase repository

Primary sequence information for precursor and mature miRNAs were procured from miRBase (version 22.1), a primary miRNA registry database for miRNA (Kozomara et al., 2019). The database contained miRNA annotated sequences from different species including *Homo sapiens*. The information was further used for deriving secondary and tertiary structure for query or prediction using different publicly available algorithms (Table 2 and Figure 2) semi-automated using python script.

**Table 2:** List of algorithms used for determining the secondary and tertiary structure of miRNAs.

| SN | Algorithm Name | Algorithm Comment | Prediction Types | Reference |
| --- | --- | --- | --- | --- |
| 1 | MXFold2 | Maximum Expected Accuracy (MEA) algorithm | 2D | (Sato et al., 2021) |
| 2 | CentroidFold | Centroid-based folding algorithm | 2D | (Sato et al., 2009) |
| 3 | RNAFold | Zuker and Stiegler algorithm | 2D | (Lorenz et al., 2011) |
| 4 | RNAComposer | SimRNA and FARFAR 2 algorithm | 2D and 3D | (Biesiada et al., 2016) |
| 5 | 3dRNA | Direct coupling analysis (DCA), k-Mean algorithm | 2D and 3D | (J. Wang et al., 2019) |

**Figure 2:**
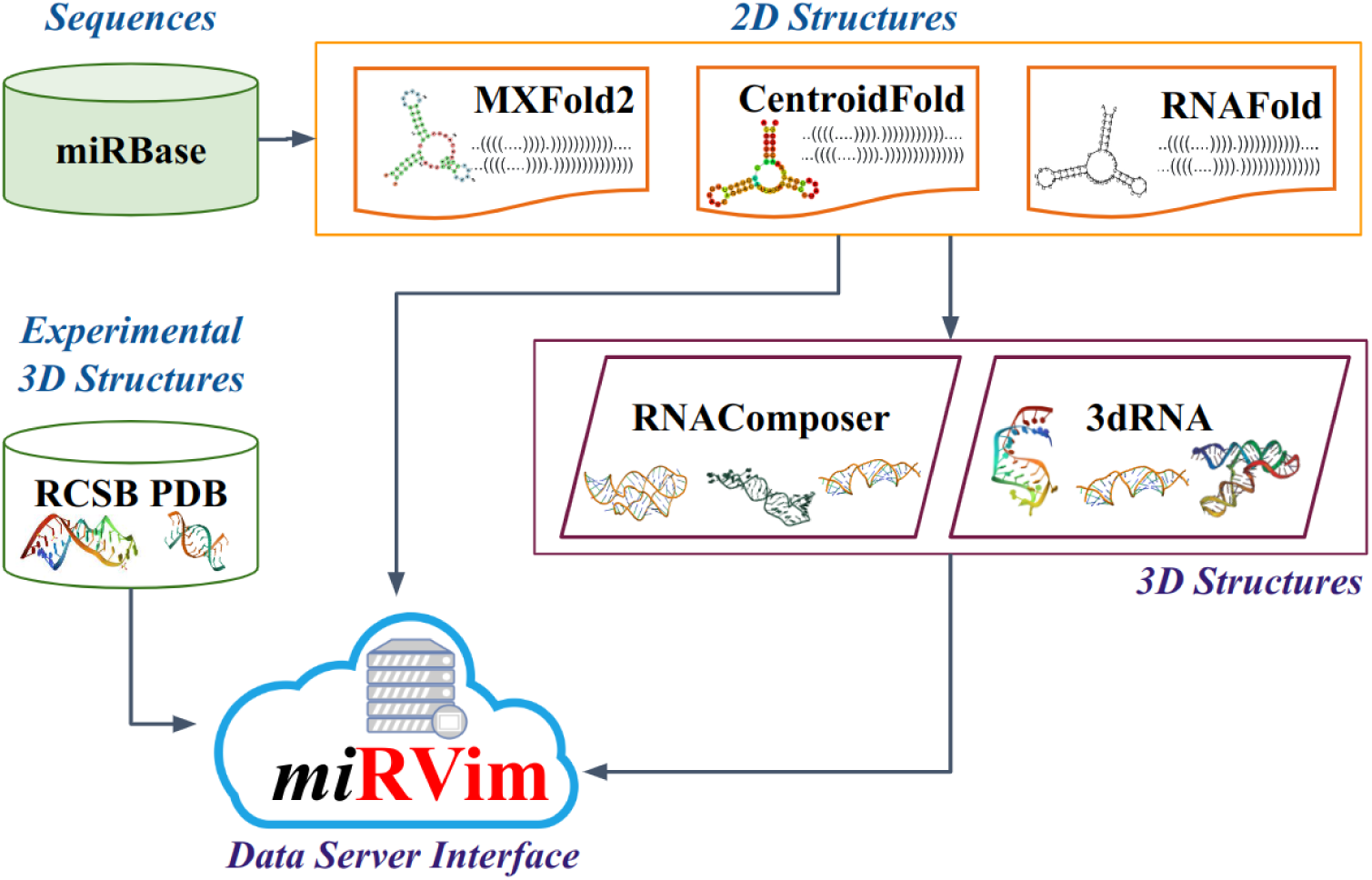
The miRVim data curation workflow to provide 3D structures from miRNA sequences

### Experimental structures of miRNAs

The Research Collaboratory for Structural Bioinformatics (RCSB) Protein Data Bank (PDB) repository was explored for experimentally determined miRNA structures (Berman et al., 2000). The repository hosts about 212 experimentally driven protein-miRNA complexes or miRNA explored by the query ‘miRNA or microRNA’ for *Homo sapiens* having multiple duplicated or partially determined structures. To get the experimentally validated miRNA structures, the sequences in FASTA format for all entries in the PDB archive file were explored for miRNA sequences and identifiers. The available structures using the selected identifiers were downloaded. The miRNA model from these structures were extracted using ChimeraX command line interface (Pettersen et al., 2021).

### 2D miRNA predicted structure curation

Three algorithms were explored to determine the two-dimensional (2D) structure of miRNAs *viz.,* MXFold2 (Sato et al., 2021), CentroidFold (Sato et al., 2009), and RNAfold (Lorenz et al., 2011) (Table 1). MXFold2 employs deep learning based methods to predict RNA secondary structures by utilising folding scores generated through a deep neural network. These scores are computed based on Turner’s nearest-neighbour free energy parameters (Turner & Mathews, 2010). The deep neural network is trained within the framework of maximum margin, incorporating thermodynamic regularisation techniques to enhance structural robustness. This regularisation ensures that the folding scores generated by the method closely align with the free energy values computed from the thermodynamic parameters. Subsequently, MXFold2 predicts the optimal secondary structure by maximising the sum of scores associated with nearest-neighbour loops, employing a dynamic programming methodology reminiscent of Zuker’s algorithm (Zuker, 2003). To train the deep neural network, MXFold2 adopts the structured support vector machine (SSVM) approach within the max-margin framework, minimising a structured hinge loss function while concurrently applying thermodynamic regularisation to minimise discrepancies between the folding scores and the thermodynamic parameter-derived free energy.

CentroidFold predicts secondary structure by employing the γ-centroid estimator, a method based on statistical decision theory (Hamada et al., 2009). It assesses the alignment between a true RNA structure and a candidate structure using a gain function that considers weighted true positives and true negatives of base pairs, with a weight factor for base pairs. The expected gain function, aiming to maximise concordance, is computed by considering various secondary structures under a given posterior distribution. This involves determining a candidate structure based on the base-pairing probability, representing the confidence in predicted base pairs and it allows the adjustment of specificity and sensitivity through the weight parameter to predict the best secondary structure.

The RNAfold primarily predicts the minimum free energy (mfe) structure for a single RNA sequence, employing the classic Zuker and Stiegler algorithm (Zuker & Stiegler, 1981). Additionally, it offers the capability to compute equilibrium base pairing probabilities using John McCaskill’s partition function algorithm (McCaskill, 1990). The output includes the predicted mfe structure represented as dot bracket notation, along with links to generated visualisation plots. These plots come in three types: firstly, a conventional secondary structure graph is generated using the naview layout method. Secondly, the pair probabilities are visualised in a “dot plot”. Lastly, a mountain plot is produced, illustrating both the predicted mfe structure and pair probabilities. This mountain plot takes the form of an xy-graph that displays the number of base pairs enclosing each sequence position. For miRVim the structures were obtained as dot-bracket notation (Mattei et al., 2014) and stored in SQL based databases for further use.

### Prediction of 3D miRNA structures

Apart from exploring databases with experimental structure availability, prediction algorithms capable of predicting 3D structure from RNA sequence were explored. Two algorithms RNAComposer (Biesiada et al., 2016) and 3dRNA (J. Wang et al., 2019) were used for the prediction of 3D miRNA structures.

RNAComposer involves building a 3D model by assembling selected 3D structural elements. Further it refines the final 3D model using CHARMM Force Field to minimise the torsion angle space and Cartesian atom coordinate space. Similarity of the input and output sequences and purine–pyrimidine compatibility, secondary structure topology compliance, and energy determine the criteria for ranking of predicted models (Biesiada et al., 2016). The RNAComposer predicted four structures and its ranking system was used as a criteria for the selection of the best model for the initial version of miRVim .

3dRNA (J. Wang et al., 2019) evolved as an automated method for generating 3D structures of RNA from provided sequences and secondary structures. It finds smallest secondary elements (SSEs) encompassing stem (helix) and loop (viz., hairpin, internal, bulge, pseudoknot, and junction) motifs. The predicted RNA structures are constructed by combining sequence data obtained from native RNA structures found in the Protein Data Bank (PDB), and secondary structure information derived using DSSR (Dissecting the Spatial Structure of RNA) using Infernal homology based search tool (Nawrocki & Eddy, 2013). In the assembly process, it assembles structures using templates from previously solved native RNA structures, templates from the same PDB ID are intentionally excluded to evaluate the overall performance of 3dRNA. The Infernal package is employed to facilitate multiple sequence alignment from the target RNA sequence. Direct Coupling Analysis (J. Wang et al., 2017) optimises RNA 3D structure prediction using evolutionary restraints of nucleotide–nucleotide interactions where only the top N values are retained, with N calculated by multiplying the RNA sequence length by a factor of 0.2. All initial structures serving as starting points for optimization are generated through the assembly procedure within 3dRNA. The algorithm incorporates simulated annealing Monte Carlo simulation, the k-Means clustering method, the Bi-residue method, and distance geometry techniques for prediction purposes. Finally, the 3dRNA scoring system (J. Wang et al., 2015) is employed to select the most suitable RNA structure.

### SASA and volume calculation

Solvent accessible surface area (SASA) calculations were performed using the FreeSASA library, an open-source C library specifically designed for SASA calculations (Mitternacht, 2016). The SASA values were obtained separately for polar and nonpolar surface areas. Additionally, the volume of the miRNA 3D structures was calculated using the ’*measure volume*’ utility in ChimeraX (Pettersen et al., 2021). This utility allows for accurate volume measurements of the biomolecular structures. The volume calculation helps in determining the overall size and spatial occupancy of the miRNA structures.

### Data processing, file management and web interface

Python and pandas library was used for data processing, data storage, and file management. Web interface was designed using front end web technologies and frameworks. 3DMol.js (Rego & Koes, 2015) for molecule visualisation, Drupal 9 for website backend framework, Codeigniter 4 for API calls, MySQL or SQLite for data storage, PetiteVue.js as javascript framework, CSS3 for designing of the visual elements of the webpage.

## Acknowledgements

Authors acknowledge Dr. D. Y. Patil Biotechnology and Bioinformatics Institute, Pune, India and Dr. D. Y. Patil Vidyapeeth, Pune, India for providing research infrastructure and opportunities. Ankita acknowledges Central University of South Bihar, Gaya, India as a host institute for providing research opportunities. Vishal Kumar Sahu acknowledges the Council of Scientific and Industrial Research (CSIR), New Delhi for CSIR-SRF fellowship (File No.: 09/1340(11487)/2021-EMR-I).

## Author Contributions

VKS and ASP curated secondary and tertiary structures of miRNAs, prepared online database & web interface, and drafted the manuscript. SB and AR planned, guided and drafted the manuscript.

